# The Distribution of Elastin and Collagen Underpinning the Smart Properties of the Interosseous Membrane

**DOI:** 10.1101/2025.01.19.633808

**Authors:** Sotiria Anastopolous, Melissa L. Knothe Tate

## Abstract

The interosseous membrane (IOM), a ligament-like structure spanning the radius and ulna, reduces strain in the ulna and structurally stiffens the radio-ulnar complex of the forearm. Using two-photon and second-harmonic-imaging we measured collagen and elastin signal intensity to test the hypothesis that their spatial distributions correspond to predominant loading patterns in the IOM. Distinct spatial gradients in collagen and elastin, as well as cruciate ligament-like architectures, were observed at the submicron and the micron to mesoscopic length scales. Quantitative analysis revealed anisotropies in the elastin-collagen composite comprising the IOM, with elastin 4-6 times higher than collagen concentrations at radius/ulna - IOM interfaces, and organized in the tensile loading direction, *i*.*e*. along the major Centroidal Axis, of the IOM. Hence, the IOM exhibits a composite structure comprising elastin and collagen, with spatial distribution of elastin higher than collagen at bone-IOM interfaces and decreasing from the interface with the ulna to that of the radius. These increased concentrations of elastin at interfaces are expected to confer elasticity (spring function). In contrast, peaks in collagen concentrations represent collagens’ organization into fibers, parallel to the length of the IOM, bridging the radius and ulna, and conferring toughness and damping function to the IOM and forearm construct. Mapping the cross-scale elastin and collagen composition of the IOM gives unprecedented insight into its emergent properties and associated mechanical function, an understanding of which may guide future surgical treatments, implant and medical textile design and manufacture, as well as physical therapy protocols to promote healing.

## Introduction

The interosseous membrane (IOM) is an essential connecting element of syndesmosis fiber joints, defined as “complex fibrous joints between two bones and connected by ligaments and a strong membrane” [1], allowing for slight movement. The respective radio-ulnar and tibio-fibular complexes exemplify syndesmosis fiber joints of the resepctive forearm and lower leg. Macroscopically, the interosseous membrane exhibits ligament-like structure (**Figure 1**). Microscopically, the IOM shares similarities with other physiological membranes with advanced functional properties, including the periosteum and tendons [2-8].

**Figure 1.**
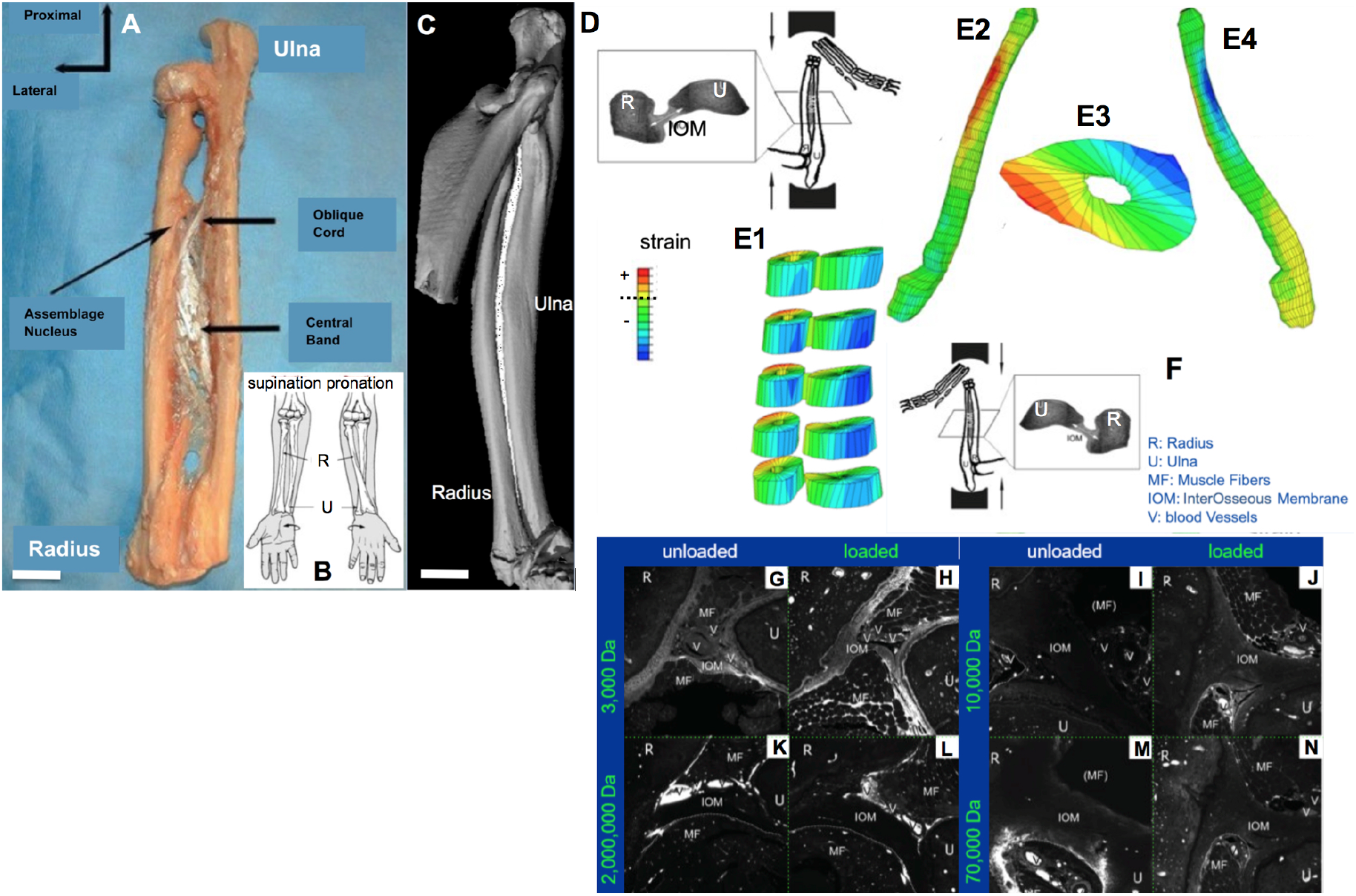
**The interosseous membrane**, (**A**) spanning the radius and ulna of the forearm in a human specimen (left, not to scale, *adapted from* [1], *used with permission*), (**B**) enables the radio-ulnar complex to achieve complex movements such as supination and pronation, [2 - *used with permission*]. (**C**) A rendering of the rat forearm, with the IOM spanning the interface between the radius and ulna, in white (not to scale, *adapted from* [7-10], *used with permission*). Of particular note, at the start of this study, the detailed anatomy of the rat IOM, *e*.*g*. cord and band structure seen in human IOM, had not been reported. The scale bar is *circa* 7 cm in the human [11] and 5 mm in the rat. **(D) The end loading model** of the rat forelimb, *after* [9,10], experimental set up (upper left quadrant, viewed from behind for comparison with finite element [FE] model) and FE model showing maximal longitudinal strain in the distal radius and ulna (**E1**,**E2**,**E4**) and transverse medio-lateral strain gradient (**E3**), *after* [7]. **(G-N) Molecular tracer studies for tracers of specific molecular weights, using the end loading model (F**, viewed from the front with structures labeled) demonstrated load-augmented molecular transport and sieving of tracers up to 70,000 Da across the tissues of the IOM and forearm complex, *after* [8]. R: Radius, U: Ulna, MF: Muscle Fibers, IOM: InterOsseous Membrane, V: blood Vessels.

The IOM is so essential to forearm function that no load transfer would occur between the radius and ulna without the IOM [2-7]. It spans between the radius and ulna and modulates the mechanics of the forearm by reducing strain on the ulna and structurally stiffening the radius-ulna complex [4]. Without the IOM, neither rotation of the forearm and hand such that the palm faces upward or forward (supination), nor downward or backward (pronation) would be possible [3]. Using finite element modeling of the rat forelimb compressive loading model, previous studies have demonstrated the role of the IOM in forelimb mechanics (**Figure 1E1-4**) [7-10]. Using this established experimental mechanobiology loading model, under axial compression, the forearm is subjected to bending loads, with a medial to lateral gradient in strain in the transverse plane (**Figure 1E3**).

If one considers the load transfer and bridging elements of the skeleton, including ligaments, tendons, the periosteum, and the interosseous membrane, there is impetus to understand the analogies and differences between the different components and how they relate to their properties as advanced functional materials [12-17]. In a series of experimental studies in combination with the aforementioned experimental rat forelimb loading model, the IOM has been shown to exhibit active barrier function, and load-modulated molecular transport and sieving, both in the IOM and across the IOM into the tissue compartments of the joint complex. Testing molecular probes from 3000 up to 2 million Daltons, loading was shown augments the transport of the molecular probes through the tissues, up to 70,000 Daltons, at which point the effect of loading is attenuated (**Figure 1G-N**). This is analogous to other tissues in which this effect has been observed, from bone to other musculoskeletal tissues [8].

In a series of imaging studies elucidating periosteum’s smart material properties, it was shown that the intrinsic patterns of elastin and collagen making up the tissue weave of periosteum underpin its mechanical-stimuli-responsive function, inspiring the design and manufacture of advanced functional materials, i.e. woven fiber composites, using recursive logic and Microscopy-Aided Design And ManufacturE (MADAME) [12-17]. This understanding is expected to open new avenues for advanced functional textiles [12-17] which lend themselves to medical, transport, safety and other applications.

At the start of the current study, our working hypothesis was that the IOM shares analogous properties with other interface tissues of the musculoskeletal system such as periosteum [12]. Specifically, we tested the hypothesis that the IOM exhibits spatial distributions of elastin and collagen specific to the major and minor axes of loading. Our approach was to study the concentration and distribution of elastin and collagen, the two predominant structural proteins making up the IOM, along the major and minor axes. Samples were analyzed in three regions, from proximal to distal, and corresponding approximately to those defined for the human IOM (**Figure 1**). For comparison, we tested the same hypothesis in the periosteum of the radius and ulna which are bridged by the IOM.

## Methods

We studied the distribution of elastin and collagen within sections of the polymethylmethacrylate (PMMA) embedded rat forearm (*n* = 5), including the ulna and radius with the IOM between, as well as adjoining muscle tissue, cut in the transverse plane at *circa* 500-micron intervals along the length of the forearm. Samples were analyzed in approximately equally divided thirds (radial | central | ulnar), centered about the line bisecting the IOM (**Figure 2**). The imaging methods were all adapted from our established, published protocols [12], described in detail below. Procion red was tracked as an established marker of blood perfusion within tissues [8,15,16]. Two photon excitation microscopy (TPIM) was used to track elastin distributions and second harmonic imaging microscopy (SHIM) for collagen. Each interosseous membrane was separated into three respective zones, and a minor and a major centroidal axis were defined to relate the distribution of collagen and elastin to predominant historical loading regimes, according to our previously published protocols [18]. Distributions of the resulting collagen and elastin signals were measured relative to the major and minor axes of each sample (Fiji, Matlab) [8]. For comparison, we measured protein fiber level to tissue fabric level differences in architectures of the periosteum, of the radius and ulna.

**Figure 2.**
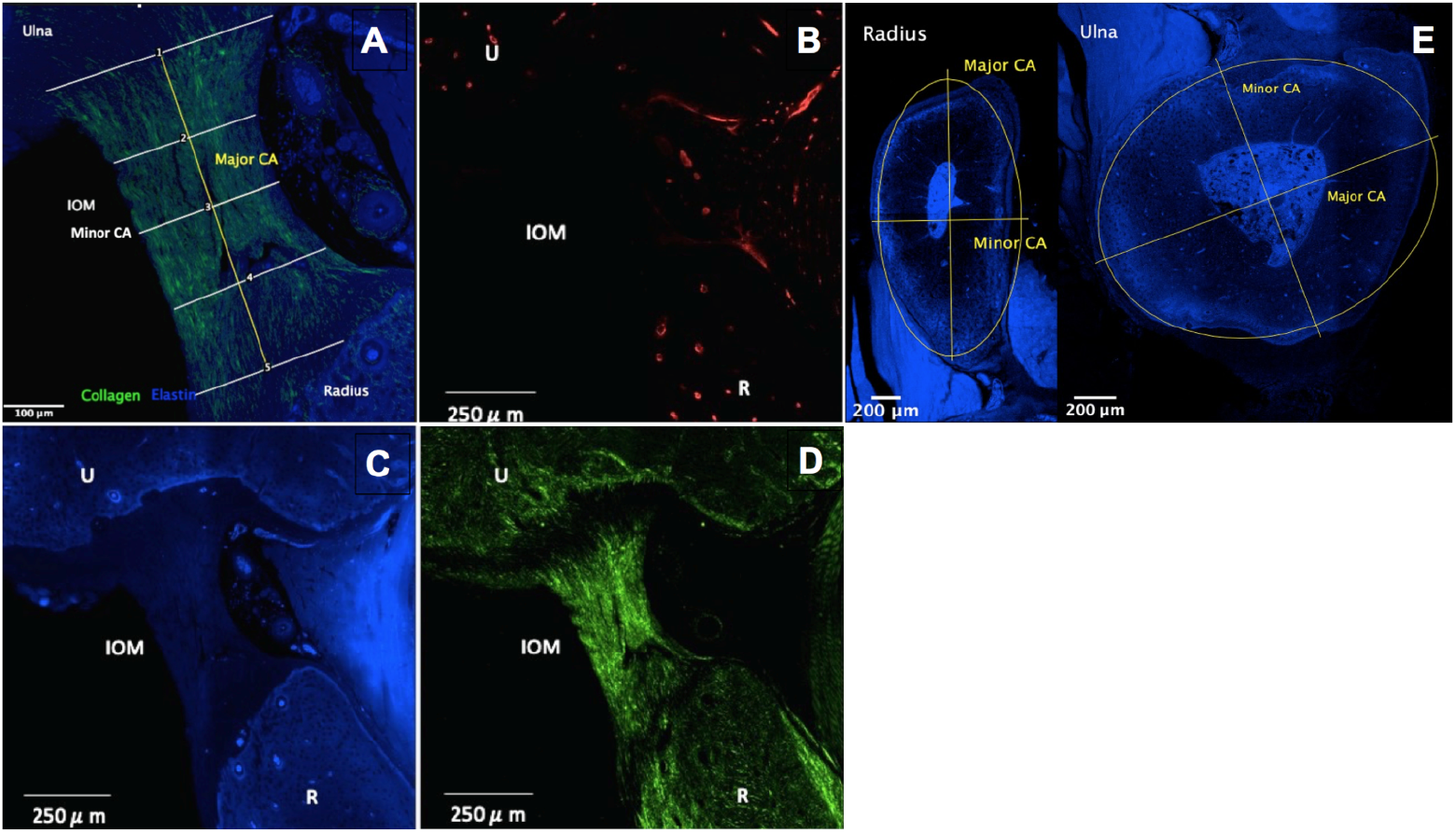
Imaging of perfusion (Procion red), elastin and collagen with respect to mechanical loading history (major and minor centroidal axes) and three regions of IOM between the radius and ulna, as well as the periosteum. **A**. To test whether distribution of elastin and collagen shows significant correlation to loading history, the major (yellow line) and minor (white line, labeled ‘3’) centroidal axes were mapped out, corresponding to the axis parallel to the radioulnar length of the IOM and the orthogonal axis, respectively. The elastin (blue, **C**.) and collagen (green, **D**.) distributions are overlaid for a typical specimen. **B**. Perfusion is tracked (red) using the intravital tracer Procion Red. **E**. Thickness of the periosteum in the radius and ulna was measured for comparison of fabric length scale correlations to loading history.

### Sample preparation

Five female Sprague Dawley rats, weighing between 215 and 240g, were anesthetized via isoflurane inhalation (0.5 - 3%) for the preterminal experiment. Body temperature was monitored and maintained at 37 deg Celsius throughout the experiment using a rectal thermometer and heating pad. All methods were carried out in accordance with the relevant guidelines and regulations of the Institutional Animal Care and Use Committee (IACUC, Protocol #98-498 Amended 09.05.2000).

To visualize and quantify bone perfusion, 0.8% Procion red solution was injected intravenously via the lateral tail vein (Imperial Chemical, London), at a dosage 0.01 ml/g body mass and 5 minutes prior to euthanasia [8,16]. Immediately after euthanasia, the forelimbs with periosteum and surrounding muscle layer *in situ* were resected and prepared for fixed, undecalcified histology using our established and standardized protocols [8]. The resulting PMMA embedded tissue blocks were sectioned transversely, every 500 µm, using a Leica diamond blade microtome. After polishing to *circa* 150 µm (Buehler Automet 2000 Polisher), sections were mounted on glass slides with glass coverslips (Eukitt).

To relate tissue fabric organization to prevalent mechanical loading histories, the major and minor centroidal axes of the bone cross sections were marked. The major and minor centroidal axes correspond respectively to the bending axes about which long bones are most (major CA, line of tension in the IOM) and least (minor CA) likely to resist a given bending load [18]. The major loading directions of the IOM were marked as parallel and orthogonal to the length of the IOM with respect to the respective ulna and radius cross sections.

### Imaging Protocol

The specimens were imaged, adapting previously established and published protocols [12], developed for imaging on a Leica SP5 II inverted microscope equipped with a Spectra Physics MaiTai HP DeepSea titanium sapphire multiphoton laser tuned to 830 nm (∼100 fs pulse), an xyz high precision multipoint positioning stage and a 63 × 1.3NA glycerol objective. The forward propagated second harmonic collagen signal was collected in the transmitted Non-Descanned-Detector using a 390–440 nm bandpass filter. The two-photon excitation of elastin was excited at 830 nm and collected in the photo-multiplier tube (PMT) using a 435–495 nm emission filter. This filter was used to segment away autofluorescence that was observed in the green channel that did not completely correspond to elastin architecture. For the Procion red signal, a 561 nm excitation was collected in the PMT using a 580–650 nm emission. A tiled scan was collected at the four quadrants correlating to the major and minor centroidal axes in the previously mentioned 3 channels, plus brightfield. Each 246 µm × 246 µm tile was imaged at 12-bit with a scan speed of 100 Hz and a resolution of 2.081 pixels per micron. Within the tiled area, a high-resolution xyz stack, with a step size of 0.5 µm and a voxel size of 0.48 µm × 0.48 µm × 0.50 µm, was acquired to capture the distribution of collagen, elastin and vasculature of the specimens in three-dimensional space.

Adaptation of the aforementioned protocol for the Zeiss LSM 880 required an 840nm excitation wavelength for both the second harmonic signal of collagen and the two-photon excitation of elastin with the former signal being detected in the transmitted Non-Descanned Detector using a 415-425nm bandpass filter and the latter signal being detected in the reflected Non-Descanned Detector using a 430-490nm bandpass filter. The Procion red signal was acquired using a 561nm excitation wavelength and collected in the PMT using a 415-735nm emission filter. A 20 × 0.8 objective (ZEISS Microscopy, Germany) was used to obtain scans of the radius and ulna.

The identified regions of interest were imaged using a 63 × 1.4 objective (ZEISS Microscopy, Germany) with Immersol immersion fluid (ZEISS Microscopy, Germany). The samples were imaged at 8-bit with a field of view comprising 224.92µm x 224.92µm and z-stacks utilized a voxel size of 0.141µm x 0.141µm x 1.000µm. The IOM of each sample was imaged using a 20 × 0.8 objective (ZEISS Microscopy, Germany). The samples were imaged at 8-bit with a field of view comprising 224.92µm x 224.92µm and z-stacks utilized a voxel size of 0.141µm x 0.141µm x 1.000µm.

With respect to signal intensity, tracer intensity can be considered as directly proportional to concentration, as *per* previous studies from our group [19] and others [20]. In this sense it can be considered as a relative comparable or as a surrogate measure for concentration. For data between the ulna and radius, signal intensity along the length was compared; for other measures pixels (signal per unit tissue area) were compared. Furthermore, potential photobleaching effects on data comparisons were mitigated by imaging all sections in an identical fashion, *i*.*e*. exposing all specimens to comparable laser light magnitude and duration.

### Quantitative Imaging and Statistical Analyses

All acquired images were processed using Image J2 (NIH Image J2 v1.49). Imaging data was filtered using Image J2 smoothing function to reduce noise. Contrast was enhanced by 0.3% adjustment in all channels. For quantification, intensities of the elastin and collagen signals were mapped from the radius to the ulna along the major and minor axes of the IOM (**Figure 7**). The **null hypothesis** that no differences in collagen and elastin concentration could be attributed to location along the major or minor CA of the IOM was tested using nonparametric Mann-Whitney test. For analysis of spatial distribution between the ulna and radius (major axis IOM), a best fit curve (red) was overlaid on raw data (Minitab 19, Minitab Inc., State College, PA). Data was then averaged for the central-most region of the IOM and the respective thirds of data collected nearest the radius and ulna.

IOM measures were compared to those of the periosteum, where in addition to measurements of elastin and collagen content in the major and minor CA of the periosteum (fiber level architecture), the thickness of the periosteum was measured (fabric level architecture). Pixels per unit area (collagen and elastin density, respectively) were quantified for elastin and collagen, in the respective major and minor axes of the periosteum (see below), to enable comparison with previously published data from the ovine femur [12]. The collagen and elastin density of the periosteum was assessed at four points on both the radius and ulna, with two points corresponding to either end of the respective bone’s major centroidal axis, and two corresponding to the endpoints of the minor centroidal axis. The value was assessed through measurement of the number of pixels counted in a selected region, divided by the area of the region to give a measurement of pixel density. Measurements at each of these identified points were carried out to a depth of 10 µm and averaged, with twelve replicates. Again, for these measures, the **null hypothesis** that no differences in collagen and elastin concentration or in could be attributed to location along the major or minor CA was tested using nonparametric Mann-Whitney test.

Periosteal thickness was measured at four points on both the radius and ulna, with two points corresponding to either end of the respective bone’s major centroidal axis, and two corresponding to the endpoints of the minor centroidal axis. Measurements at each of these identified points were carried out to a depth of 10µm, resulting in a sample size of n=10 for each test point. The nonparametric Mann-Whitney test was used to test the **null hypothesis** that no difference in thickness of the periosteum could be attributed to location on the minor or major centroidal axis.

Statistical analyses were carried out using Minitab 19 and Microsoft Excel where significance was defined at a p value of 0.05 (95% confidence interval), using Mann-Whitney testing of null hypotheses, since the data were not normally distributed.

## Results

### Qualitative Observations of the IOM

From proximal to distal, along the IOM, the structures observed in the human forelimb IOM (**Figure 1**) are observable in the rat forelimb IOM. The oblique cord of the IOM appears remarkably similar to the cruciate ligament of the knee (**Figure 3**). The collagen fibrils align parallel to the oblique cord, fanning out at insertion points to the ulna and radius. The Procion signal is quite strong in the vein (diffuse red, left vein below IOM) and artery (below and to the right of the vein, with red blood cells fluorescing with Procion), as well as in the muscle tissue (top right, bottom left quadrants with fascicles fluorescing).

**Figure 3.**
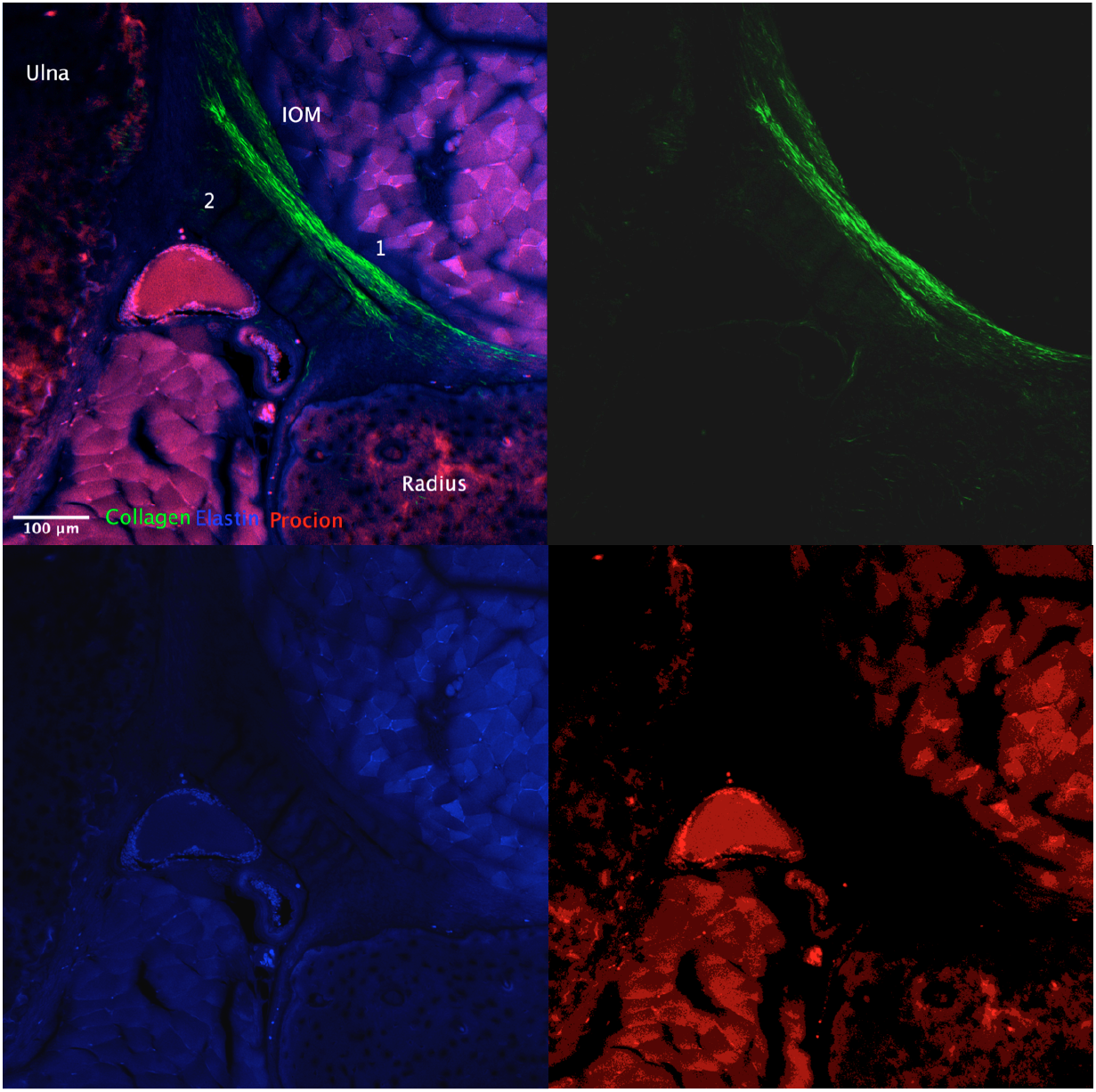
High resolution imaging of collagen and elastin distribution as well as vascular perfusion (Procion red) in the oblique chord, at the proximal segment of the IOM and surrounding tissues. In this set of images (clockwise), three separate channels corresponding to the channel overlay (top left), collagen (top right), elastin (bottom left), and Procion (bottom right) channels.

Moving proximally down to the middle section of the IOM, there is more of a band structure with remarkable changes in the patterns of elastin and collagen distribution at points of insertion in the ulna and in the radius. The strong presence of Procion red indicates perfusion in blood vessels and the lack of blood vessels within the IOM (**Figure 4**).

**Figure 4.**
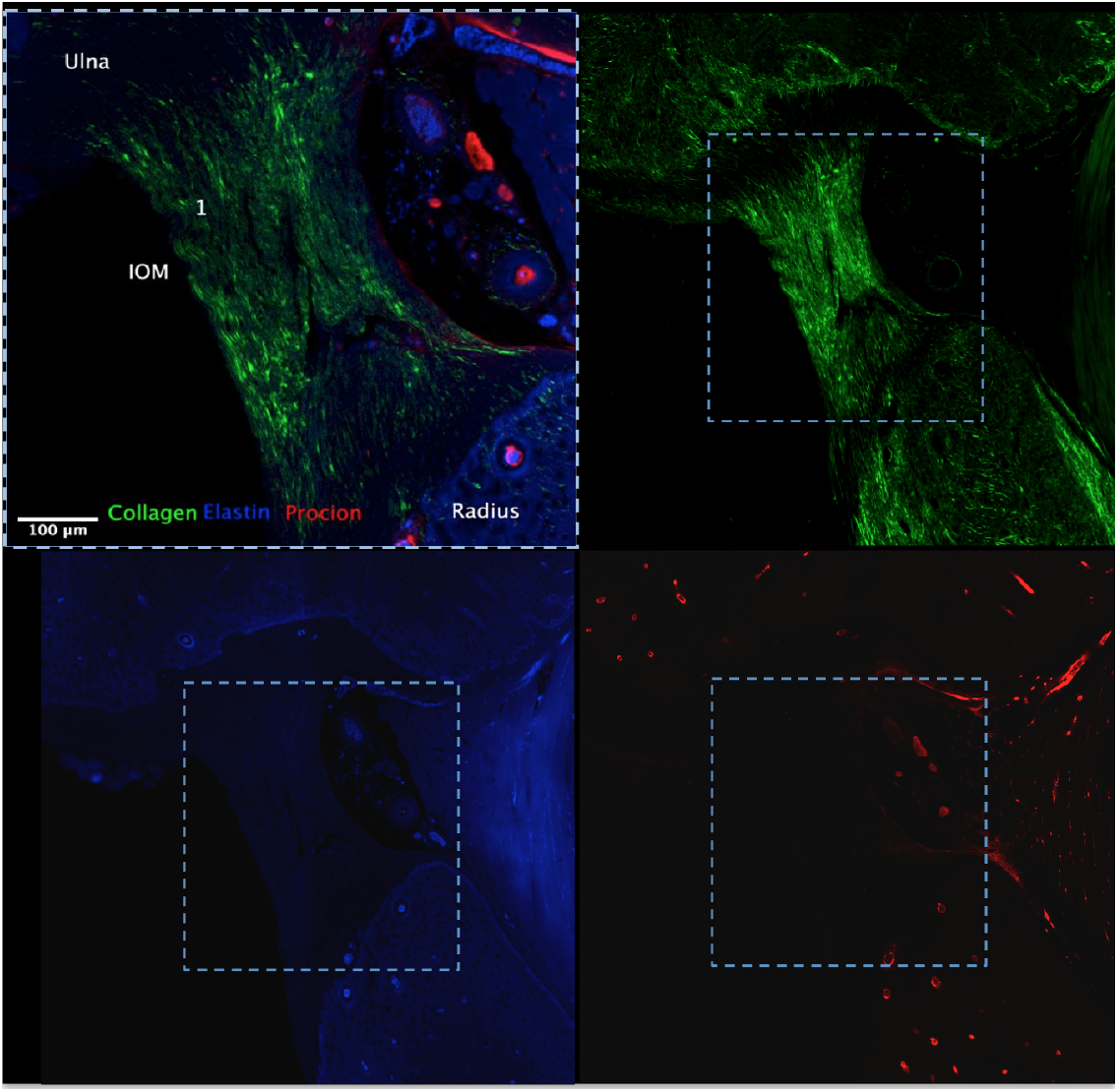
High resolution imaging of collagen and elastin distribution as well as vascular perfusion (Procion red) in the central band of the IOM and surrounding tissues. In this set of images (clockwise), three separate channels corresponding to the channel overlay (top left, area depicted by the dashed square in subsequent images), collagen (top right), elastin (bottom left), and Procion (bottom right).

In the distal most portion (**Figure 5**) this representative image shows the remarkable change in the structure where the IOM is essentially interwoven with the elastin. The IOM surrounds the blood vessel structure and inserts into the radius. Finally, the high-resolution images enable visualization of the IOM’s myriad insertions into the bone of the radius (**Figure 6**). They look remarkably like Sharpey’s fibers, which are prevalent in the periosteum, at insertion points into bone. Similar to Sharpey’s fibers of periosteum, the IOM fibers insert into the surface of the radius at approximately 45-degree angles.

**Figure 5.**
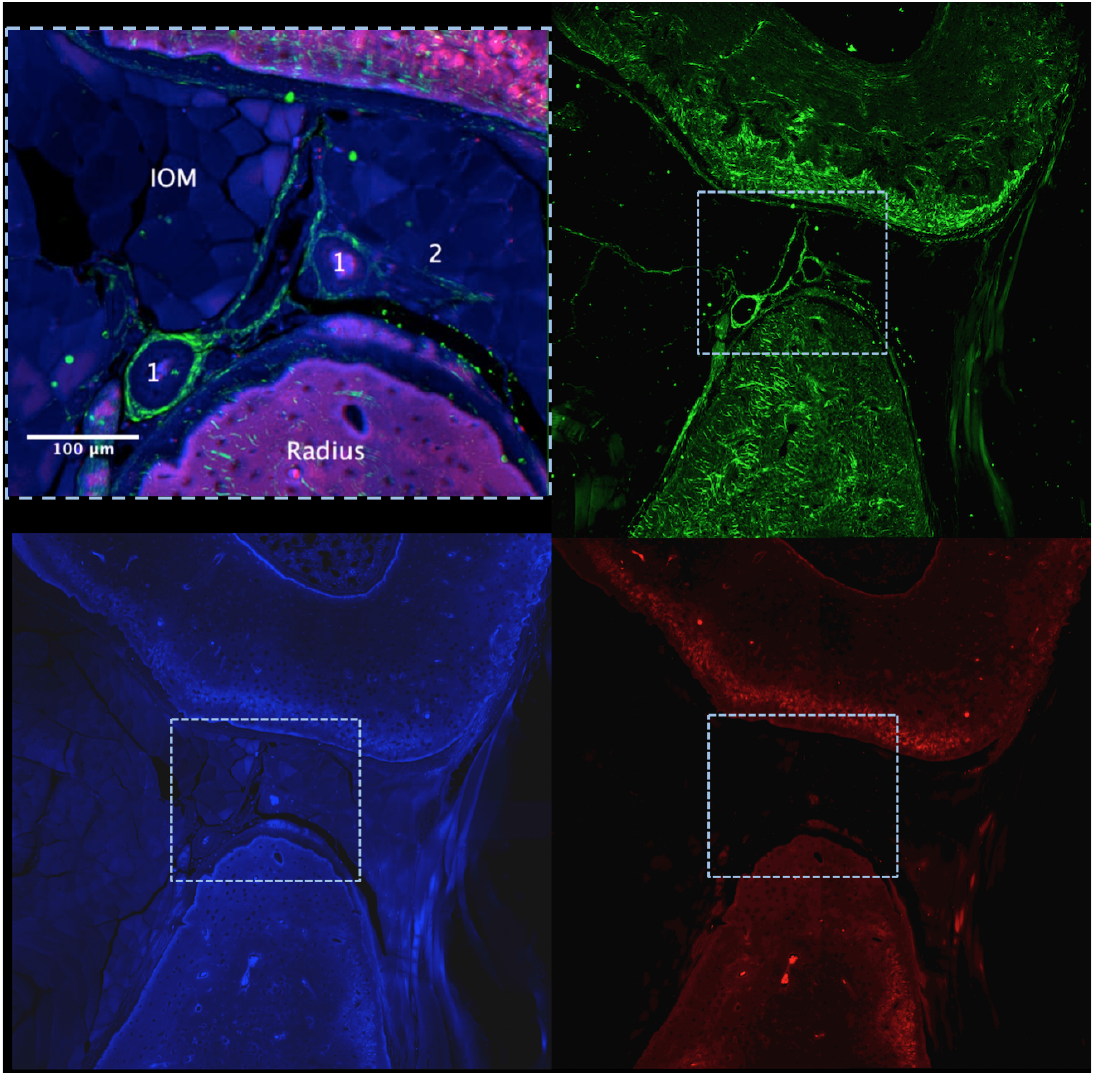
High resolution imaging of collagen and elastin distribution as well as vascular perfusion (Procion red) on the distal edge of the IOM and surrounding tissues. In this set of images (clockwise), three separate channels corresponding to the channel overlay (top left, area depicted by the dashed square in subsequent images), collagen (top right), elastin (bottom left), and Procion (bottom right).

**Figure 6.**
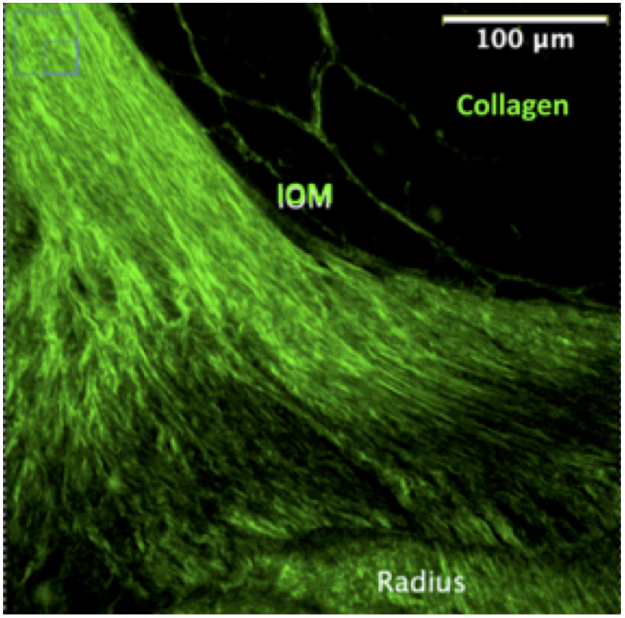
**High resolution imaging of collagen distribution** where the IOM inserts into the radius, with Sharpey’s fibers inserting at 45 degree angles at the Radius.

### Hypothesis Testing Based on Quantitative Measures of the IOM

We first measured the signal intensity of the respective elastin and collagen signals quantitatively as signal intensity, from the radius to the ulna, *i*.*e*. along the major centroidal axis. This is a measure for relative concentration, as all samples are probed using the same microscope and laser excitation and emission settings [7,8,12]. No significant differences or trends were observed in the amounts of collagen or elastin with respect to the minor axis of the IOM (data not shown). In contrast, along the major axis of the IOM, for five representative animals (*n* = 5) with at least three samples per animal, similar trends were observed in the distribution of both collagen and elastin, as well as between the radius and ulna. Namely, concentrations of elastin and collagen paralleled each other, following roughly similar slopes (rates of slight increase or decrease) along the length of the IOM (**Figure 7**) Based on the best fit curve (red) (overlaid on raw data, Minitab 19, Minitab Inc., State College, PA), mean values for the radial, central and ulnar segments along the major centroidal axis showed a significant decrease in elastin, with a small but significant increase in collagen going from radial to ulnar sections of the IOM. Hence, the **null hypothesis** that no differences in collagen and elastin concentration could be attributed to location was confirmed along the minor CA of the IOM but was negated for the major CA.

**Figure 7.**
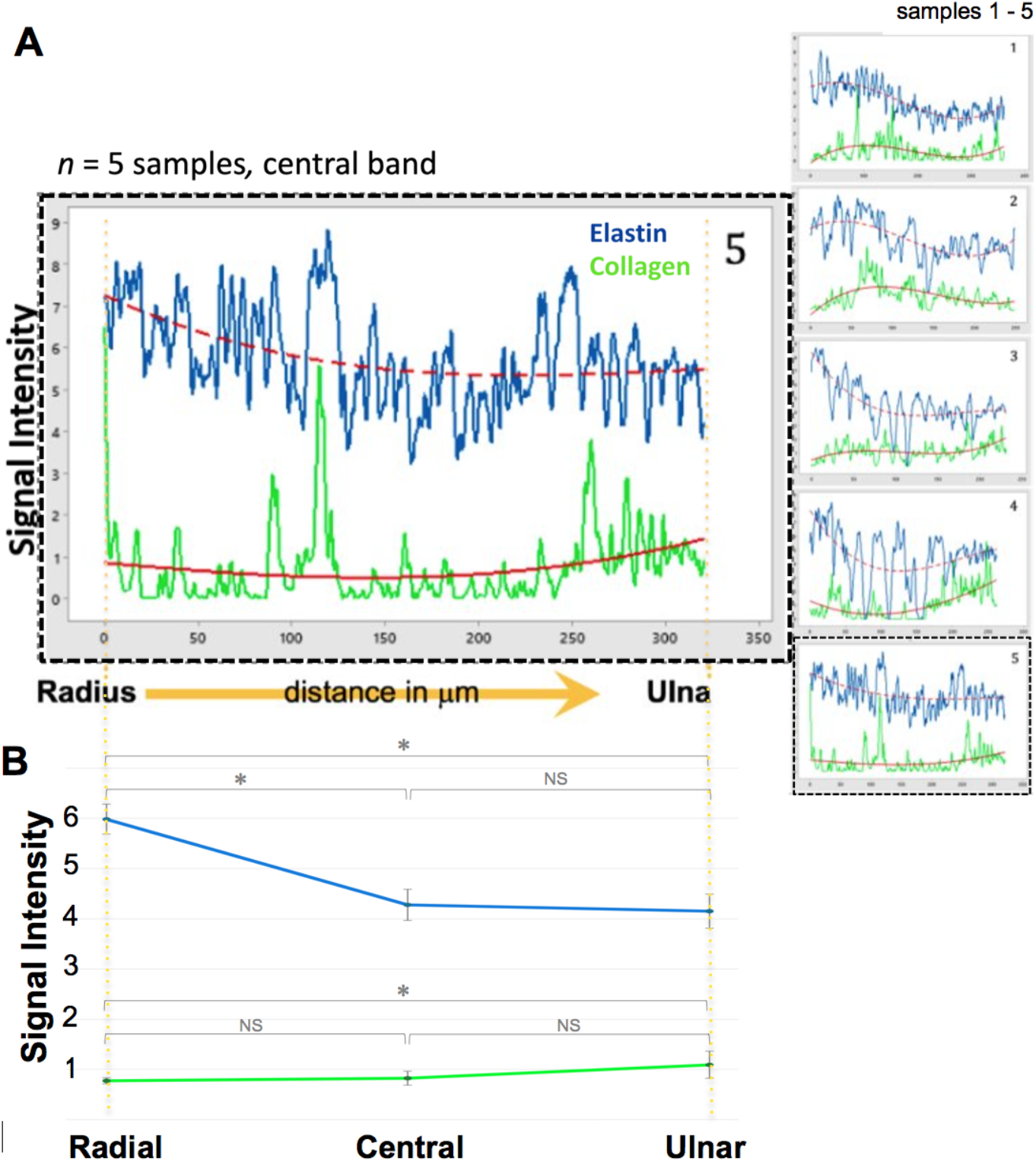
**Quantitative assessment of the elastin and collagen signal along the length of the IOM**, from the radius (left) to the ulna (right), with best fit curves (red) overlaid on raw data. **A**. Signal intensity for five forelimb samples, with sample 5 highlighted. Signal intensity is a relative surrogate measure for concentration [7,8,12], as all microscope and laser excitation and emission settings are held constant throughout the measurements. Intensities increase and decrease slightly, roughly in parallel along the distance of the central band of the IOM, from the interface with the radius to the ulna, respectively, measured in microns. **B**. Mean signal intensity +/-standard error of the mean for the radial, central and ulnar third of the IOM, as measured in A. Significant differences are observable in elastin intensity between the radial and central and radial and ulnar band of the IOM, and in collagen intensity between the radial and ulnar band of the IOM. *Significance defined as p < 0.05, using ANOVA, with Nonsignificant differences indicated by NS, *n =* 5 samples.

Of note, micro to mesoscopic patterns of the elastin and collagen emerged as architectures when viewed at the mesoscopic length scale (**Figures 3-6**). In general, the IOM exhibits a composite structure comprising elastin and collagen. The spatial distribution of elastin decreases along the length of the IOM, with increasing proximity to the ulna, and its concentration (as measured by intensity) remains more than four times higher than that of collagen along the length of the IOM (going from six times higher at the radial edge down to four times higher at the ulnar edge, **Figure 7B**).

### Quantitative Measurements of the Periosteum

Measurements of signal density (pixels per unit area) for elastin and collagen at the major and minor centroidal axes of the periosteum, for the respective radius and ulna cross sections (**Supplementary Table 1, Supplementary Figure 1**), showed no significant differences attributable to loading direction, *e*.*g*. along the major or minor axis, the anterior or posterior aspect of the bone, or the bone itself (radius *versus* ulna), thus confirming the null hypothesis. In contrast, periosteal thickness measurements showed significant differences attributable to presence within the radius or ulna, respectively, the anterior or posterior radius or ulna, and the major and minor centroidal axis of the radius; no significant differences were measured in periosteum thickness between the major versus minor centroidal axis of the ulna (**Table 1, Figure 8**), thus confirming the null hypothesis regarding differences related to mechanical loading (minor and major CA) of the ulna but negating the same hypothesis specific to the radius, and negating the null hypothesis regarding differences intrinsic to the radius and ulna irrespective of mechanical loading function.

**Table 1.** Median periosteal thickness, null hypothesis testing using the Mann-Whitney test. In all but two cases (shaded cells), significant differences (defined by *p < 0.05) were observed, rejecting the null hypothesis.

| Sample Site | Median ($\mu\text{m}$) | p value |
| --- | --- | --- |
| Ulna | 25.3 | 0.001 |
| Radius | 29.9 |  |
| Ulna Radius major CA | 28.7 | 0.718 |
| Ulna Radius minor CA | 29.7 |  |
| Ulna major CA | 26.5 | 0.337 |
| Ulna minor CA | 23.9 |  |
| Radius major CA | 36.3 | 0.000 |
| Radius minor CA | 29.7 |  |
| Ulna Radius Posterior | 22.7 | 0.000 |
| Ulna Radius Anterior | 29.9 |  |
| Ulna Posterior | 22.7 | 0.009 |
| Ulna Anterior | 26.4 |  |
| Radius Posterior | 22.3 | 0.000 |
| Radius Anterior | 36.6 |  |

**Figure 8.**
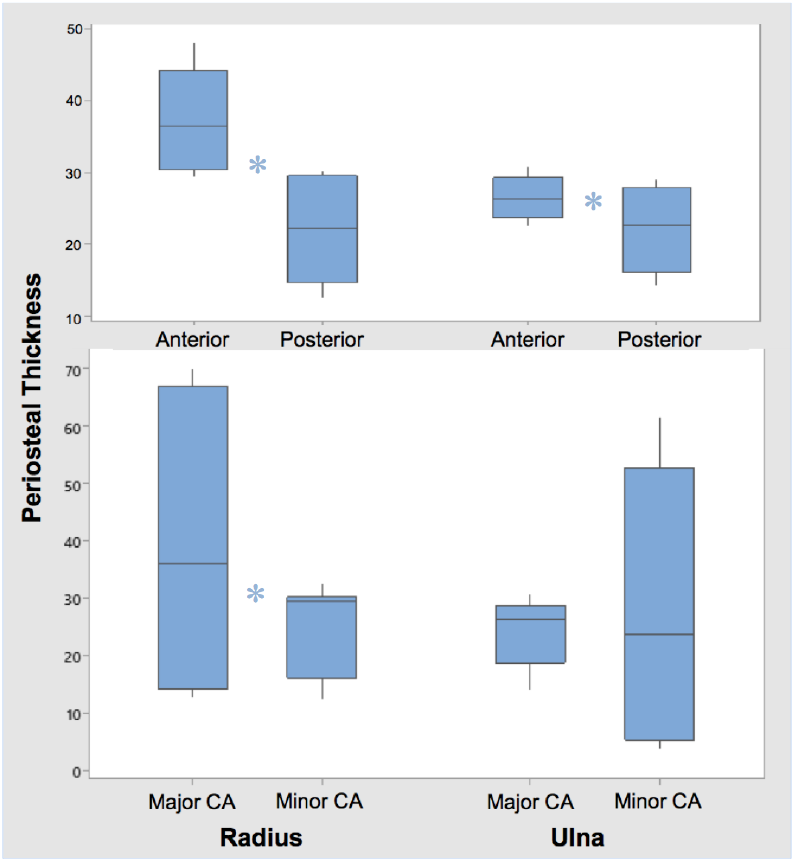
Graphical depiction of hypothesis testing using the Mann-Whitney test, where significant differences between groups are indicated by *p < 0.05.

## Discussion

Hypothesis testing demonstrated significant effects of structural protein (*i*.*e*. elastin and collagen) concentration (intensity) along the length of the IOM, which corresponds to the major CA, *i*.*e*. the predominant direction of tensile loading. Such effects were not observed along the minor CA which is least resistant to loading. In contrast, no significant differences in collagen and elastin density (area equivalent to concentration along a line) were observed in association with the major and minor CAs of the periosteum of the respective radius and ulna. Interestingly, significant differences were observed in the thickness of the periosteum in association with predominant loading directions (major/minor CA) in the radius but not in the ulna, as well as in association with anatomic markers, *i*.*e*. posterior *versus* anterior, of both the radius and ulna. Taken together these quantitative observations have interesting implications for multi-length scale structure-function relationships in band-like (IOM, ligament, tendon) *versus* beam-like (long bone) composites, where predominant loading direction and function is expected to exert greater influence on tissue fabric patterns comprising concentrations and distributions of structural proteins imbuing elasticity and toughness [12,13], *i*.*e*. where interfaces likely dictate both structure and function. In the case of composite beam structures comprising bone, both the loading environment and material properties likely exhibit additional complexity as described in further detail below.

The IOM exhibits a composite structure comprising elastin and collagen, with spatial distribution of elastin relatively constant albeit approximately four to six times higher than collagen at bone-IOM interfaces. Approximately fifty percent (and significantly) more elastin was observed in the section of the IOM nearest the radius than in the central or section of IOM nearest the ulna, corresponding to bone interfaces under highest strain under compression (**Figure 1**); increased concentrations of elastin at interfaces is expected to confer elasticity (spring function). In contrast, peaks in collagen concentrations represent collagens’ organization into fibers, parallel to the length of the IOM, bridging the radius and ulna, and with a small but significant increase near the IOM interface with the ulna; this is expected to confer toughness and damping function to the IOM and forearm construct.

The relative concentrations of elastin:collagen are similar in the IOM as those reported previously in periosteum of the ovine femur, i.e. 1:0.25 or four times more elastin [12]. Direct comparisons could not be made with the radius or ulna of the current study samples, as no statistically significant differences were observed in their relative concentrations of elastin:collagen, which may in itself reflect on structure:function relationships of the IOM and periosteum in syndesmosis fiber joints compared to long bones of the forearm, respectively, hindlimb. Specifically, the periosteum has been shown previously to contribute to the superior ultimate strength, stiffness, absorbed energy and deflection of intact rat femora when loaded in three-point-bending to failure [23], as compared to periosteum-stripped femora. In addition, the periosteum of the ovine femur has been shown previously to exhibit intrinsic prestress *in situ* when an abundance of Sharpey’s fibers “velcro” it to the femur [24,25]. In addition, the ovine femur periosteum exhibits anisotropic stiffness in the axial compared to circumferential direction [26] and anisotropic permeability in the bone to muscle and muscle to bone directions [24]. These respective smart properties of ovine periosteum tissue reflect the respective distibution of both elastic (elastin) and tough (collagen) structural protein fibers and tight junctions between cells in the inner, cellular cambium layer of the periosteum [24].

As noted above, it is interesting to consider biomechanical implications of a soft-tissue sleeve (periosteum) “velcroed” onto the surface a hindlimb long bone such as the femur compared to a soft-tissue band of the IOM joining the ulna and radius of the forearm syndesmosis fiber joint. While periosteum tissue circumferentially surrounds long bones like a sleeve, the IOM exhibits a band-like structure between the nearest roughly linear surfaces of two adjoining bones, e.g. in this case the radius and ulna. The spatial distributions of collagen and elastin in both the periosteum of the femur as well as the periosteum and IOM spanning the ulna and radius appear to be influenced by boundary conditions as well as prevailing, tensile loading modes, which likely in turn relate to tissue growth prior to skeletal maturity and postnatal healing throughout life [27]. In the case of the IOM, the major Centroidal Axis aligns with a band of tissue exposed to repeated cyclic tension under compressive loading of the forelimb [7,8].

These data demonstrate a previously underappreciated aspect of musculoskeletal structure and function, providing impetus for further imaging, across and between length scales, to elucidate the interaction of structure-function relationships, from the molecular to the organ and organ system scale, among different tissues of the musculoskeletal system. These experimentally determined parameters provide critical data for mechanical modeling and subsequent understanding and prediction of the IOM’s mechanical function as well as replacement of that structure/function after trauma or disease.

## Conclusions

In addition to its mesoscale organization into cord and band structures spanning the ulna and radius, the IOM’s compositional weave of collagen and elastin play an important role in conferring toughness and elasticity, which in turn modulates the IOM’s mechanical function. Mapping the cross-scale elastin and collagen composition of the IOM gives unprecedented insight into its emergent properties and associated mechanical function, an understanding of which may guide future surgical treatments, medical textile and implant designs, and physical therapy protocols to promote healing [12-17,21,22]. For example, gradient weaves emulating the distribution of elastin and collagen at insertion points to the ulna and radius may provide new insights to address current hurdles associated with tendon and ligament repair. While stress concentrations can be mitigated by such gradients if the natural tissue weaves develop during growth of the organism (with a multitude of insertion points developing *in situ* and adapting with use over time), seamless gradients are currently challenging to replicate e.g. after tendon and ligament failure. Similarly, recent approaches to emulate structural protein architectures of the periosteum are finding use in design and manufacture of wearable sleeves (e.g. compression garments) and implantable (orthopaedic, vascular stent) devices.

---

**Supplementary Table 1.** Median collagen and elastin densities for null hypothesis testing using Mann-Whitney Test. No significant differences (defined by p < 0.05) were observed, confirming the null hypothesis.

| Sample Site | Median<br>Pixels/$\mu\text{m}^2$ | p value |
| --- | --- | --- |
| **Collagen Density** |  |  |
| Ulna | 48.3 | 0.977 |
| Radius | 49.2 |  |
| Ulna major CA | 38.7 | 0.173 |
| Ulna minor CA | 49.3 |  |
| Radius major CA | 46.7 | 0.936 |
| Radius minor CA | 49.2 |  |
| Ulna Radius Posterior | 48.9 | 0.471 |
| Ulna Radius Anterior | 39.1 |  |
| **Elastin Density** |  |  |
| Ulna | 48.0 | 0.844 |
| Radius | 49.3 |  |
| Ulna major CA |  | 0.21 |
| Ulna minor CA | 28.9 |  |
|  | 48.0 |  |
| Radius major CA | 43.9 | 0.835 |
| Radius minor CA | 49.3 |  |
| Ulna Radius Posterior |  | 0.173 |
| Ulna Radius Anterior | 22.8 |  |
|  | 4.95 |  |

**Supplementary Figure 1.**
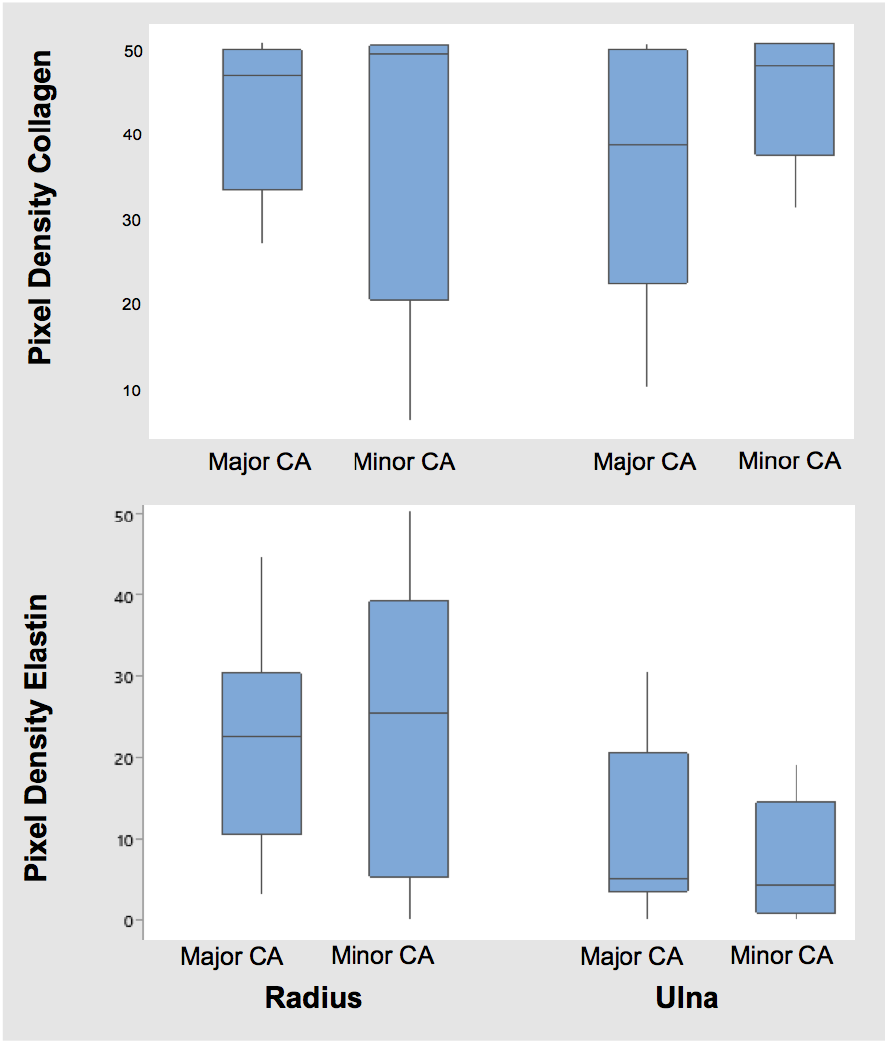
Graphical depiction of hypothesis testing using the Mann-Whitney test, where significant differences between groups are indicated by *p < 0.05. No significant differences were observed, thus supporting the null hypothesis.

## References

[1] physio-pedia.com, last accessed on 5 May 2024.

[2] Dumontier C, Soubeyrand M (2014) The forearm joint, In: Bentley G (Editor) European Surgical Orthopaedics and Traumatology, Springer, Berlin.

[3] Birkbeck DP, Failla JM, Howshaw SJ, Fyhrie DP, Schaffler M (1997) The interosseous membrane affects load distribution in the forearm, Journal of Hand Surgery, 22(6): P975–980.

[4] Rougereau G, Marty-Diloy T, Vigan M, Vialle R, Soubeyrand M, Langlais T (2022) Biomechanical assessment of the central band of the interosseous membrane using shear wave elastography: reliability and reproducibility, Journal of Hand Surgery-European Volume, 47(11): 1134–1141, doi:10.1177/17531934221114301

[5] Schneiderman G, Meldrum RD, Bloebaum RD, Tarr R, Sarmiento A (1993) The Interosseous Membrane of the Forearm-Structure and Its Role in Galeazzi Fractures, Journal of Trauma, 35(6): 879–885.

[6] Kholinne E, Kwak J-M, Sun Y, Koh KH, Jeon I-H (2021) Forearm Interosseous Ligaments: Anatomical and Histological Analysis of the Proximal,Central, and Distal Bands, Journal of Hand Surgery-American Volume, 46(11):29, doi: https://doi.org/10/1016/j.jhsa.2021.03.002

[7] Tami AE, Suresh G, Patel RB, Knothe Tate ML (2004) Effect of Interosseous Membrane on Load Transfer in Rat Forelimb using Finite Element Analysis, Proceedings of IMECE04, 2004 ASME International Engineering Congress BED TOC: IMECE2004-61934: 181–182.

[8] Knothe Tate ML, Tami A, Netrebko P, Milz S, Docheva D (2012) Multiscale computational and experimental approaches to elucidate musculoskeletal and ligament mechanobiology using the ulna-radius-interosseous membrane construct as a model system, Tech Health Care, 20(5):363–78. doi: 10.3233/THC-2012-0686

[9] Uthgenannt B, Silva MJ (2007) Use of the rat forelimb compression model to create discrete levels of bone damage in vivo, J Biomech, 40(2): 317–24, doi: 10.1016/J.JBIOMECH.2006.01.005

[10] Robling AG, Hinant FM, Burr DB, Turner CH (2002) Improved bone structure and strength after long-term mechanical loading is greatest if loading is separated into short bouts, J Bone Min Res, 17: 1545–1554.

[11] Carrillo F, Suter S, Casari FA, Sutter R, Nagy L, Snedeker JG, Fürnstahl P (2020) Digitalization of the IOM: A comprehensive cadaveric study for obtaining three-dimensional models and morphological properties of the forearm’s interosseous membrane, Sci Reports 10: 6401, 10.1038/s41598-020-63436-3

[12] Ng J, Knothe L, Whan R, Knothe U, Knothe Tate ML (2017) Scale-up of Nature’s Tissue Weaving Algorithms to Engineer Advanced Functional Materials, Nature Sci Reports, 7: 40396(1-10), doi: 10.1038/srep40396

[13] Knothe Tate ML (2020) Advanced design and manufacture of mechanoactive materials - inspired by skin, bones and skin-on-bones. Frontiers in Bioengineering and Biomaterials|Biomaterials 8: 845(1-11), doi: 10.3389/fbio.2020.00845

[14] Anastopolous S, Ngo L, Ng JL, Putra, V.D.L., Knothe Tate ML (2024) Interface tissues of the mesoderm: periosteum, ligament, interosseous membrane, and myofascial tissues, an inspiration for next generation medical textiles. Current Opinion in Biomedical Engineering,

[15] Knothe Tate ML, Dolejs S, McBride S, Miller RM, Knothe UR (2011) Multiscale Mechanobiology of De Novo Bone Generation, Remodelling & Adaptation of Autograft in a Common Ovine Femur Model, Journal of the Mechanical Behavior of Biomedical Materials 4(6): 829–40. doi: 10.1016/j.jmbbm.2011.03.009

[16] Ngo L, Knothe Tate ML (2023) A spike in circulating inflammatory cytokines alters barrier function between vascular and musculoskeletal tissues, Nature Scientific Reports, 13, 9119; 10.1038/s41598-023-30322-7.

[17] Sidler HJ, Duvenage J, Anderson EJ, Ng J, Hageman DJ, Knothe Tate ML (2018) Prospective design, rapid prototyping and testing of smart dressings, drug delivery patches, and replacement body parts using Microscopy Aided Design And ManufacturE (MADAME), Frontiers in Medicine | Translational Medicine 5: 348(1-11), 10.3389/fmed.2018.00348

[18] McBride SH, Dolejs S, Miller RM, Knothe U, Knothe Tate ML (2011) Major and Minor Centroidal Axes Serve as Objective, Automatable Reference Points to Test Mechanobiological Hypotheses Using Histomorphometry, Journal of Biomechanics 44(6):1205–1208. doi: 10.1016/j.jbiomech.2011.01.033

[19] Ngo L, Knothe Tate ML. A spike in circulating cytokines TNF-α and TGF-β alters barrier function between vascular and musculoskeletal tissues. Sci Rep. 2023 Jun 5;13(1):9119. doi: 10.1038/s41598-023-30322-7.

[20] Lee K, Shirshin E, Rovnyagina N, Yaya F, Boujja Z, Priezzhev A, Wagner C. Dextran adsorption onto red blood cells revisited: Single cell quantification by laser tweezers combined with microfluidics. Biomed. Opt. Express. 2018;9:2755. doi: 10.1364/BOE.9.002755.

[21] Rosetti L, Küntz LA, Kunold E, Schock J, Müller KW, Grabmayr H, Stolberg-Stolberg J, Pfeiffer F, Sieber S-A, Burgkart R, Bausch AR (2017) The microstructure and micromechanics of the tendon-bone insertion, Nat Mater 16: 664–670, doi: 10.1038/nmat4863.

[22] Deymier AC, An Y, Boyde JJ, Schwartz AG, Birman V, Genin GM, Thomopolous S, Barber AH (2017) Micro-mechanical properties of the tendon-to-bone attachment, Acta Biomater 56: 25–35, doi: 10.1016/j.actbio.2017.01.037.

[23] Yiannakopoulos CK, Kanellopoulos AD, Trovas GP, Dontas IA, Lyritis GP (2008) The biomechanical capacity of the periosteum in intact long bones, Arch Orthop Trauma Surg 128:117–20. doi: 10.1007/s00402-007-0433-5.

[24] Evans SF, Parent JB, Lasko CE, Zhen X, Knothe UR, Lemaire T, Knothe Tate ML. Periosteum, bone’s “smart” bounding membrane, exhibits direction-dependent permeability. J Bone Miner Res. 2013 Mar;28(3):608–17. doi: 10.1002/jbmr.1777.

[25] Evans SF, Chang H, Knothe Tate ML. Elucidating multiscale periosteal mechanobiology: a key to unlocking the smart properties and regenerative capacity of the periosteum? Tissue Eng Part B Rev. 2013 Apr;19(2):147–59. doi: 10.1089/ten.TEB.2012.0216.

[26] McBride SH, Evans SF, Knothe Tate ML (2011) Anisotropic mechanical properties of ovine femoral periosteum and the effects of cryopreservation, J Biomech 44:1954–9. doi: 10.1016/j.jbiomech.2011.04.036.

[27] Moore SR, Milz S, Knothe Tate ML (2013) Periosteal thickness and cellularity in mid-diaphyseal cross-sections from human femora and tibiae of aged donors, J Anatomy 224: 142–149.

